# Targeting dendritic cell-specific TNFR2 improves skin and joint inflammation by inhibiting IL-12/ IFN-γ pathways in a mouse model of psoriatic arthritis

**DOI:** 10.1101/2024.06.20.598545

**Authors:** Raminderjit Kaur, Jennifer M. Harvey, Roberta Brambilla, Unnikrishnan M. Chandrasekharan, M. Elaine Husni

**Affiliations:** Department of Cardiovascular and Metabolic Sciences, Lerner Research Institute; The Miami Project to Cure Paralysis, Department of Neurological Surgery, University of Miami Miller School of Medicine, Miami, FL, USA; Department of Rheumatic and Immunologic Diseases, Cleveland Clinic, 9500 Euclid Avenue, Cleveland, OH 44195; Department of Biochemistry, School of Medicine, Case Western Reserve University, Cleveland, OH 44106

**Keywords:** Tumor necrosis factor-α (TNF) receptor 2, Psoriatic arthritis, Dendritic cells, Th1 cells, TNF inhibition

## Abstract

Psoriasis (PsO) and Psoriatic arthritis (PsA) are immune-mediated inflammatory diseases affecting the skin and joints. Approximately, 30% of patients with PsO develop PsA over time with both conditions being associated with elevated tumor necrosis factor-alpha (TNF-α) expression. TNF-α mediates its effect through two membrane receptors, TNFR1 and TNFR2. While current TNF-α-neutralizing agents, targeting both TNFR1 and TNFR2 receptors, constitute the primary treatment for psoriatic diseases, their long-term use is limited due to an increase in opportunistic infections, tuberculosis reactivation and malignancies likely attributed to TNFR1 inactivation.

Recent findings suggest a pivotal role of TNFR2 in psoriatic disease, as evidenced by its amelioration in global TNFR2-knockout (TNFR2KO) mice, but not in TNFR1KO mice. The diminished disease phenotype in TNFR2KO mice is accompanied by a decrease in DC populations. However, the specific contribution of TNFR2 in dendritic cells (DCs) remains unclear. Here, utilizing a mannan-oligosaccharide (MOS)-induced PsA model, we demonstrate a significant reduction in PsA-like skin scaling and joint inflammation in dendritic cell-specific TNFR2 knockout mice (DC-TNFR2KO). Notably, MOS treatment in control mice (TNFR2 fl/fl) led to an increase in conventional type 1 dendritic cells (cDC1) population in the spleen, a response inhibited in DC-TNFR2KO mice. Furthermore, DC-TNFR2KO mice exhibited reduced levels of interleukin-12 (IL-12), a Th1 cell activator, as well as diminished Th1 cells, and interferon-gamma (IFN-γ) levels in the serum compared to controls following MOS stimulation.

In summary, our study provides compelling evidence supporting the role of TNFR2 in promoting PsA-like inflammation through cDC1/Th1 activation pathways.

## INTRODUCTION

Psoriatic arthritis (PsA) is a persistent inflammatory disorder that primarily impacts the joints (1–4) and surrounding tendons known as enthesitis which leads to substantial functional limitations and diminished quality of life. PsA, if untreated, results in irreversible bone damage, in the affected joints (5–8) as well as significant decrease in health related quality of life measures. Currently, the first line treatment regimen for PsA is the use of TNF-α neutralizing agents (9–11). Unfortunately, up to 40% of patients do not respond to this treatment, and assessing its efficacy can take several months (13). Furthermore, the long-term use of these agents is linked to potentially life-threatening adverse effects. The major adverse effects associated with anti-TNF therapy include opportunistic infections, tuberculosis reactivation and certain malignancies (9,10,13). Interestingly, preclinical studies on mice have shown that inhibition of TNFR1 rather than TNFR2 inhibition leads to these adverse effects (14–16). In contrast, TNFR2 induces immune cell activation/proliferation and neo angiogenesis, known characteristics of psoriatic diseases (17,18).

Dendritic cells (DC) are a heterogeneous subset of antigen-presenting cells that links the innate and adaptive arms of the immune system. DC has been implicated in numerous inflammatory and autoimmune diseases, including PsA (19). Activated DC secrete TNF-α, IFN-γ, IL-12, and IL-23 following stimulation of pathogen/damage-associated molecular pattern receptors (21). These cytokines facilitate the differentiation of naïve T cells to the Th1 and Th17 subtypes which are crucial mediators of psoriatic diseases. Our previous investigations demonstrated that global TNFR2 depletion has a mitigating effect on imiquimod (IMQ) induced psoriasis like inflammation in murine models. Notably, the IMQ treatment elicited a response in DC subsets; such as plasmacytoid DCs, myeloid DC, and Tip-DCs, that was attenuated in the global TNFR2KO mice (20). Additionally, the decrease in DC product, IL-23, which is critical for down-stream polarization and stabilization of inflammatory Th17 cells (21), was observed in the TNFR2KO mice compared to controls in the IMQ mouse model. This elucidation underscores the potential role of TNFR2-dependent DC expansion/activation in driving the pathogenesis of psoriatic diseases.

In the present study we investigate the contribution of DC-specific TNFR2 to the pathogenesis of psoriatic disease. To test this, we compared the development of PsA-like inflammation in DC-specific TNFR2KO mice versus TNFR2-intact mice (control), utilizing a mannan-oligosaccharide (MOS) induced model (22). Our study reveals that DC-TNFR2 plays a significant role in promoting PsA-like skin lesions and joint inflammation likely through augmentation of both the abundance and function of a specific subtype of dendritic cells known as conventional dendritic cell type 1 (cDC1).

## MATERIAL METHOD

### Animals and genotyping

The animal-related protocols were approved by an institutional ethical committee. All mice used were maintained on a C57BL/6 genetic background, kept under specific pathogenic-free conditions, and provided with food and water ad libitum at the Biological Resource Unit facility. Experiments were performed on 8 to 10 weeks-old gender-matched mice. TNFR2-floxed mice (TNFR2 fl/fl) were crossed with CD11c-Cre transgenic mice (Jackson Laboratory, Strain #008068) to delete TNFR2 from dendritic cells (TNFR2 fl/fl/ CD11c-Cre) mice. TNFR2 fl/fl were used as a control. TNFR2 fl/fl mice were obtained from Dr. Roberta Brambilla (The Miami Project to Cure Paralysis, Dept. Neurological Surgery, and the University of Miami Miller School of Medicine, FL 33136, USA). Genotyping was performed on chromosomal DNA isolated from toe clips. The genotyping to detect the loxP-flanked (floxed) TNFR2 transgenic animals was performed using the following primers: 5’ TTGGGTCTAGAGGTGGCGCAGC-3’ and 5’-GGCCAGGAAGTGGGTTACTTTAGGGC-3’ resulting in products of 410 bp for wildtype and 578bp for the floxed allele. The following primers were used to detect the transgenic Cre expression: 5’-ACT TGG CAG CTG TCT CCA AG’-3’, 5’-GCG AAC ATC TTC AGG TTC TG-3’, 5-CAA ATG TTG CTT GTC TGG TG-3’ and 5’-GTC AGT CGA GTG CAC AGT TT-3’ generating products 313 bp for Cre allele and 200bp as internal positive control.

### Mannan Oligosaccharide (MOS)-induced PsA model in mice

MOS was purchased from Sigma-Aldrich Inc. USA and dissolved in PBS. Mice were given a single injection of MOS (IP, 800mg/kg body weight) on day 0. The ear and paw thickness were measured on alternative days using the electronic digital dial thickness gauge (0-0.4“/10mm caliper, 0.01mm resolution, ±0.03 mm accuracy). Mice were scored for inflammation in peripheral joints on alternative days as described previously (22). The Psoriasis Area and Severity Index (PASI) graded the severity of psoriatic skin lesions, which comprises the parameters of skin erythema, scaling, and thickness. PASI was scored on a scale from 0 to 4 in a blinded fashion. The mice of each group were PASI scored on day 0, day 2, day 4, and day 6 days. The grip strength was measured using a grip strength meter (Columbus Instruments, USA).

### Histology Evaluation of skin and joints

Paws were decalcified in TCA (A11156.30, Thermo Fisher Scientific, USA) prior to fixing and embedding. Decalcified paw and skin tissues were fixed in 10% neutral buffered formalin (5705, Epredia, USA) and embedded in paraffin. Tissue sections were stained with H& E and/or safranin O and epidermal thickness, leukocyte infiltration, and cartilage content were assessed.

### Flow cytometry analysis

CD11c+ cells from spleen underwent flow cytometry analysis. To ensure optimal recovery and purity of CD11c+ cells, spleen was initially subjected to enzymatic digestion using a spleen dissociation kit (130-095-926, Miltenyi Biotec, USA). The red blood cells were lysed using ACK lysing buffer (23). The Cd11c+ cells from the dissociated spleen preparation were isolated through magnetic separation using a negative selection kit (480098, Biolegend, USA).

The cells were then counted and stained with Live/ dead fixable dye (L34960, ThermoFisher Scientific, USA). After incubation with Fc block solution (550270, BD Biosciences, Franklin Lakes, NJ) samples were stained with extracellular antigen-specific antibodies, for CD11c, CD11b, MHCII, CD8, CD45R, CD80, CD86, and LY6C for 30 minutes at 4°C in the dark. For intracellular iNOS staining, the cells were fixed (by BD cytofix), permeabilized (by BD perm), prior to antibody treatment for 30 minutes at room temperature. The cells were then washed and analyzed using BD LSRFortessa (BD Biosciences, Franklin Lakes, NJ). The workstation is managed by FACSDiva software, version 10 (Tree Star, Ashland, OR). The data were analyzed using FlowJo v 10.8.1 (BD, 385 Williamson Way Ashland, OR 97520, USA).

To measure Th1 and Th17 cells, spleen cells were prepared as above and processed as per the user manual of the Mouse Th1/Th2/Th17 Phenotyping Kit (560758, BD Biosciences, Franklin Lakes, NJ). A cocktail of antibodies reacting to CD4, IL-17A, IFN-γ, and IL-4, was incubated for 30 minutes at room temperature in the dark. Following washing steps, cells were analyzed using BD LSRFortessa.

### Cytokine analysis

Mice was sacrificed by cardiac puncture on day 6 post-MOS administration and blood was collected in 1.5 ml Eppendorf tubes containing heparin. Following centrifugation plasma was collected and stored at −80°C before cytokine analysis by ELISA. ELISA kits for IL-12p70 (BMS6004) aIFN-*γ* (BMS609) and TNF-α (BMS607-3) were purchased from Invitrogen, thermos fisher Scientific USA.

### Statistical analysis

Statistical analyses were performed using SPSS version 28.0 (IBM, SPSS Inc. Chicago, USA) and Prism (GraphPad Software, San Diego, CA). Data are expressed as mean ± S.E and were assessed using Student’s t-test. A significance level of p < 0.05 was considered statistically significant. In figures, asterisks indicate the level of statistical significance (*p < 0.05, **p < 0.01, ***p < 0.001, and ****p < 0.0001).

## RESULTS

### PsA-like skin scaling and joint inflammation is reduced in DC-TNFR2KO

DC-TNFR2KO mice were generated by crossing TNFR2-floxed mice with CD11c-Cre mice, resulting in TNFR2 depletion in DC, confirmed via FACS analysis (Supplementary Fig.1). Notably, TNFR2-intact control mice (TNFR2 fl/fl) (Figure 1A, upper panel) exhibited visible scaling, thickness, and redness in response to MOS, which was less prominent in DC-TNFR2KO mice (Figure 1A, lower panel). The cumulative PASI score comprises of erythema + scaling + skin thickness was significantly lower in DC-TNFR2KO mice at day 4 and day 6 post-MOS treatment compared to control mice (p < 0.01, Figure 1B). Moreover, MOS–induced redness, swelling, and inflammation in the paws were attenuated in the DC-TNFR2KO mice (Figure 1C). Similarly, the Mean arthritis severity score (average numerical representation of the severity of arthritis symptoms in terms of paw swelling and erythema) significantly decreased in DC-TNFR2KO mice on day 4 (p < 0.01) and day 6 (p < 0.001) as shown in Figure 1D. Further, the loss of grip strength in the control mice due to MOS treatment was partially restored in DC-TNFR2KO mice by day 6 (p < 0.05, Figure 1E).

**Figure 1.**
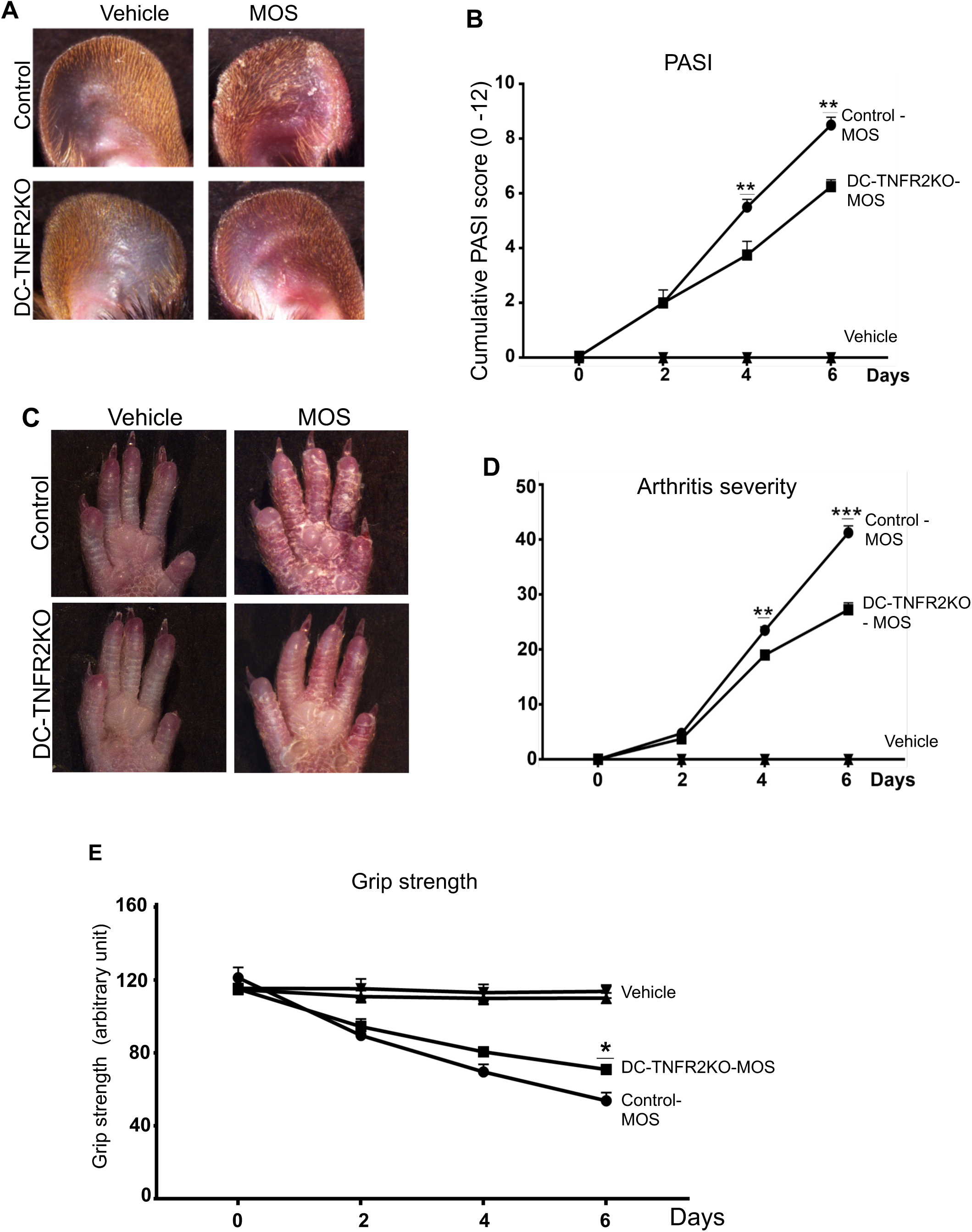
DC-TNFR2KO mice show a reduced pathological phenotype in mannan-oligosaccharide (MOS)-induced PsA model. **(A)** Representative images of vehicle or MOS treated ear of TNFR2 fl/fl (control) and DC-TNFR2KO mice. **(B)** Cumulative PASI score (erythema + scaling + thickness) of the ear by estimated in a blinded fashion. X-axis: before MOS injection (0 day) and day 2, day 4 and day 6 post-MOS injection. (**C)** Representative images of the arthritic phenotype (swelling and erythema of the paws) and psoriasis-like skin lesions in the hind paws of control and DC-TNFR2KO mice. (**D)** Mean arthritis severity score of paws and (**E**) Grip strength of front paws on indicated days (n = 6, means ± S.E, **p < 0.01; ***p < 0.001).

Histology analysis, followed by image quantification revealed a robust increase in epidermal thickness in control mice with MOS treatment. In contrast, MOS induced epidermal thickness was significantly reduced in DC-TNFR2KO mice compared to control mice (p < 0.001, Figure 2A & 2B). Furthermore, leukocyte infiltration in response to MOS was significantly diminished in DC-TNFR2KO mice compared to control mice (p < 0.01, Figure 2C). Safranin O staining indicated reduced cartilage destruction in digits of hind limp paws of DC-TNFR2KO mice relative to control mice (p < 0.05, Figure 2D & 2E).

**Figure 2.**
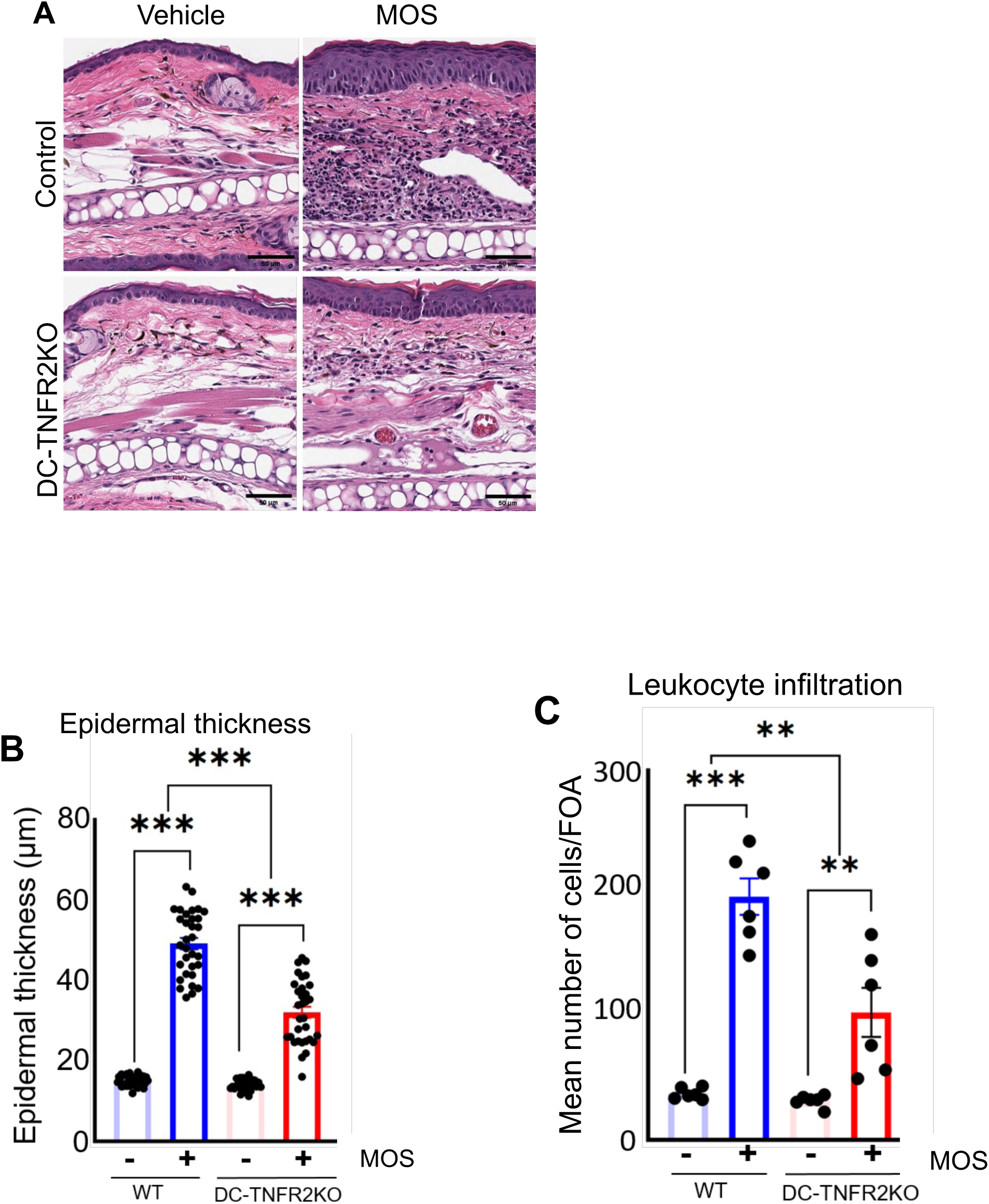
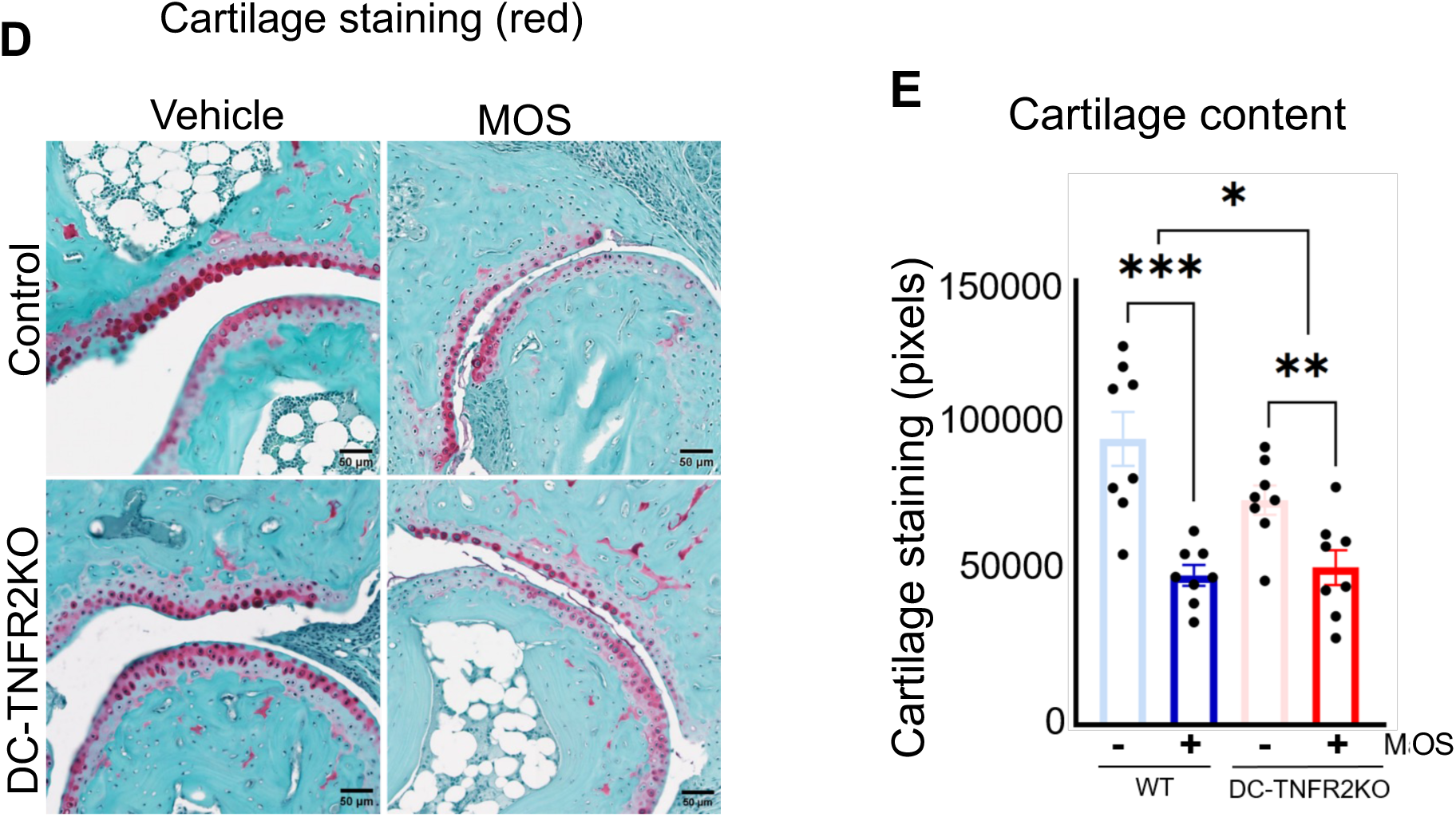
MOS-induced epidermal thickness, leukocyte infiltration and cartilage loss were diminished in DC-TNFR2KO mice. **(A)** Representative H&E staining of ear tissue from TNFR2 fl/fl (control) and DC-TNFR2KO mice ± MOS. **(B)** Epidermal thickness and, **(C)** Leukocyte count in control and DC-TNFR2KO mice ± MOS. (**D)** Representative figures of safranin O stained cartilage (red) of distal interphalangeal joints of control and DC-TNFR2KO mice ± MOS**. (E)** Cartilage content in the interphalangeal joints in control and DC-TNFR2KO mice ± MOS (n = 4, means ± S.E, * p < 0.05, p < 0.01, *** p < 0.001). Readings are taken on tissue samples harvested on day 6 post-MOS treatment.

### TNFR2 depletion attenuated the increase of conventional DC1 (cDC1) population in mice with PsA-like inflammation

Following MOS treatment, a significant increase in cDC1 population in the spleen was observed in the control mice (7.3 ± 1.55% to 14.06 ± 2.2 %, p < 0.001), whereas this increase was absent in DC-TNFR2KO mice (Figure 3A&B). The difference in cDC1 population between control and DC-TNFR2KO post-MOS treatment was significant (p < 0.01, Figure 3B). Conversely, cDC2 population did not change significantly in response to MOS in either group (Figure. 3C). Although there was an upward trend in pDC population in both control and DC-TNFR2KO mice with MOS, it did not reach statistical significance (Supplementary Fig. 2). The FACS strategy for measuring cDC1, cDC2 and pDC is shown in Supplementary Fig. 3.

**Figure 3.**
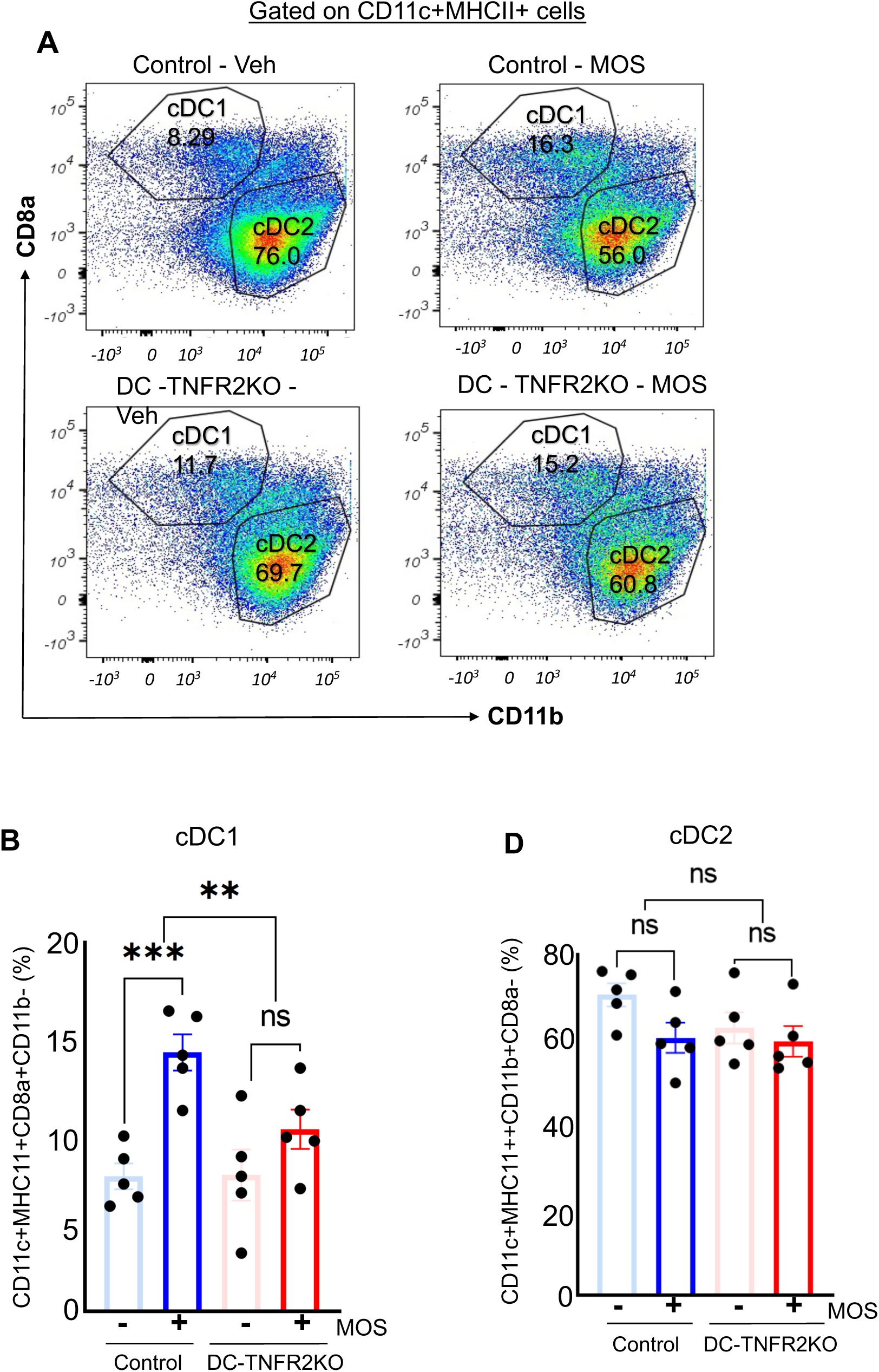
Increase in conventional DC1 (cDC1) population in the spleen following MOS treatment is reduced in DC-TNFR2KO mice. **(A)** representative figure showing the percentage of cDC1 and cDC2 cells. **(B)** Percentage of cDC1 and, **(C)** cDC2 in control and DC-TNFR2KO mice ± MOS (n = 5, means ± S.E, ** p < 0.01, *** p < 0.001). Readings are taken on tissue samples harvested on day 6 post MOS treatment.

### DC-TNFR2 is responsible for the increase of Th1 cells in PsA-like inflammation

The activated DC can induce polarization of naïve Th cells into effector T cells including Th1 and Th17 cells. Proinflammatory cytokines IFN-γ, and IL-17 produced by Th1 and Th17 cells, respectively, play a crucial in the pathogenesis of psoriatic disease (24). We observed a significant increase in Th1 cell population in the spleen of control mice following MOS treatment (p < 0.001, Figure 4A & B). In contrast, the magnitude of increase in Th1 cell population was minimal in DC-TNFR2KO mice. The difference in fold increase of Th1 cells post-MOS treatment between control and DC-TNFR2KO mice was also significant (p < 0.01, Figure 4C). Th17 cell population increased in both groups post-MOS treatment, albeit not significantly. Moreover, the levels of IL-12, IFN-γ, and TNF-α, were significantly elevated in the serum of control mice, compared to DC-TNFR2KO mice upon MOS treatment (Figures 4D-F).

**Figure 4.**
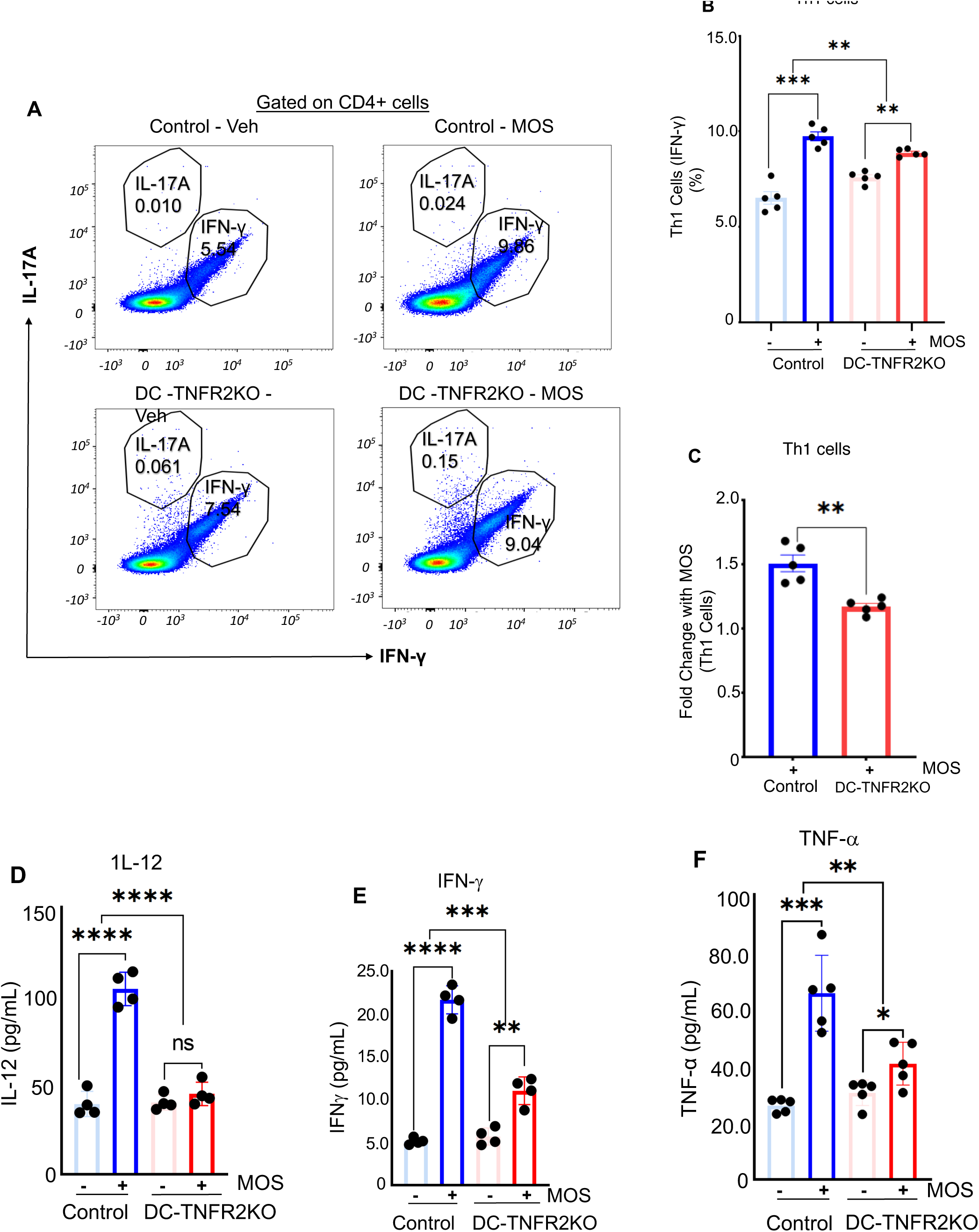
Increase of Th1 cell populations and circulating IL-12, IFN-γ and TNF-α following MOS treatment are reduced in DC-TNFR2KO mice. **(A)** Representative FACS figure showing the percentage of IL-17A^high^ (Th17 cells), IFN-γ ^high^(Th1 cells) on CD4+ gated cells. **(B)** Percentage of Th1 cells (IFN-γ ^high^) in control and DC-TNFR2KO mice following MOS treatment. **(C)** Fold change of Th1 cells in control vs DC-TNFR2KO following MOS treatment (A-C, n = 5, means ± S.E). Serum levels of **(D)** IL-12, **(E)** IFN-γ and, (**F)** TNF-α (measured by ELISA) in the control and DC-TNFR2KO mice ± MOS (D-F, n = 4, means ± S.E), * p < 0.05 **< 0.01, ***p < 0.001, ****p < 0.0001). Readings are taken on tissue samples harvested on day 6 post MOS treatment.

## DISCUSSION

Although various biological agents are recently emerged to treat PsO, anti-TNF agents remain the first-line of treatment for PsA (12). These agents neutralize TNF-α thus prevents activation of both TNFR1 and TNFR2 (25). However, the adverse effects associated with long-term use of anti-TNF agents are primarily attributed to TNFR1 inhibition (9,10,13,25). Our results demonstrating TNFR2 inactivation in DCs effectively reduces PsA-like inflammation carry significant clinical implications. This suggests that targeting TNFR2 or it’s downstream signaling pathways responsible for DC function may offer a more targeted therapeutic option compared to anti-TNF drugs for treating PsA.

Among the central players in the immune landscape, dendritic cells (DCs) are the most effective antigen-presenting cells, bridging the gap between innate and adaptive immune responses (26–28). In psoriatic diseases TNF-α increases IL-23 production in DCs, which, in turn, promote the Th17 cell polarization and down-stream effector molecules in skin and joints (29). Our study found that absence of TNFR2 in DCs significantly reduced skin and joint inflammation in the murine model of PsA induced by MOS. Although there was a notable decrease in cartilage erosion in the paw joints of DC-TNFR2KO mice compared to control mice, the overall joint phenotype was mild in this mouse model. This was expected as MOS induces robust joint disease only when reactive oxygen species (ROS) are depleted (22). We intentionally avoided macrophage overresponse in our study, as it could mask the role of the dendritic cells in disease pathology, especially considering that CD11c is expressed in macrophages and therefore CD11c-Cre can delete TNFR2 in macrophages (30,31). Further cell-specificTNFR2-depletion are needed to decipher the role of TNFR2 expressed in other immune cells contributing to PsA pathogenesis.

TNFR2 induces immune cell proliferation and activation (32), particularly, TNFR2’s role in immune suppressive T regulatory cells is well studied (33). TNFR2 can also promote proliferation of pathogenic T cells, including CD4+ and CD8+ T cells in immune-mediated inflammatory diseases (34, 35). However, role of TNFR2 in DC function or DC expansion, particularly in the case of cDCs during pathological conditions is not clear. One study, implicated TNFR2’s involvement in human cDC2 maturation in the lung, but not cDC1 maturation in response to adjuvants, in mice (36). Our study revealed that TNFR2 activity is crucial for cDC1 expansion, but not cDC2 expansion in the murine model of PsA. The cDC1 are primarily responsible for priming CD8+ T cells by cross-presentation of antigens in conjunction with MHC class I, but cDC1 also can activate Th1 cells (37). Both CD8+T and Th1 cells and can promote psoriatic diseases (36). Since mouse and human cDC1 are functionally similar (27) targeting cDC function or inducing tolerogenicity in cDC1 could be a novel approach to mitigating psoriatic disease.

In summary, this study provides previously uncharacterized role of TNFR2/cDC1-axis in promoting pathogenesis in PsA. Our findings suggest that inactivating TNFR2 while preserving TNFR1, which mediates the host-defense function of TNF-α, may ameliorate PsA effectively with potentially fewer adverse effects compared to anti-TNF agents.

## Data Availability Statement

All relevant data for this article is included, and any further inquiry including the protocol used can be made to the corresponding authors.

## Author Contributions

M.E.H, U.MC and R.K conceptualized and designed the experimental plan, analyzed the data, and wrote the manuscript. R.K and J.H performed the experiments, R.B. provided technical help and valuable reagents and provided critical discussions.

## Acknowledgement.

This work is supported by NIH Grant R01AR075777 to M.E.H & U.M.C. National Psoriasis Foundation (NPF) Bridge-Grant: NPF1808UC to U.M.C. & M.E.H. Early Research Grant from National Psoriasis Foundation to R.K.

---

**Supplementary Figure 1.**
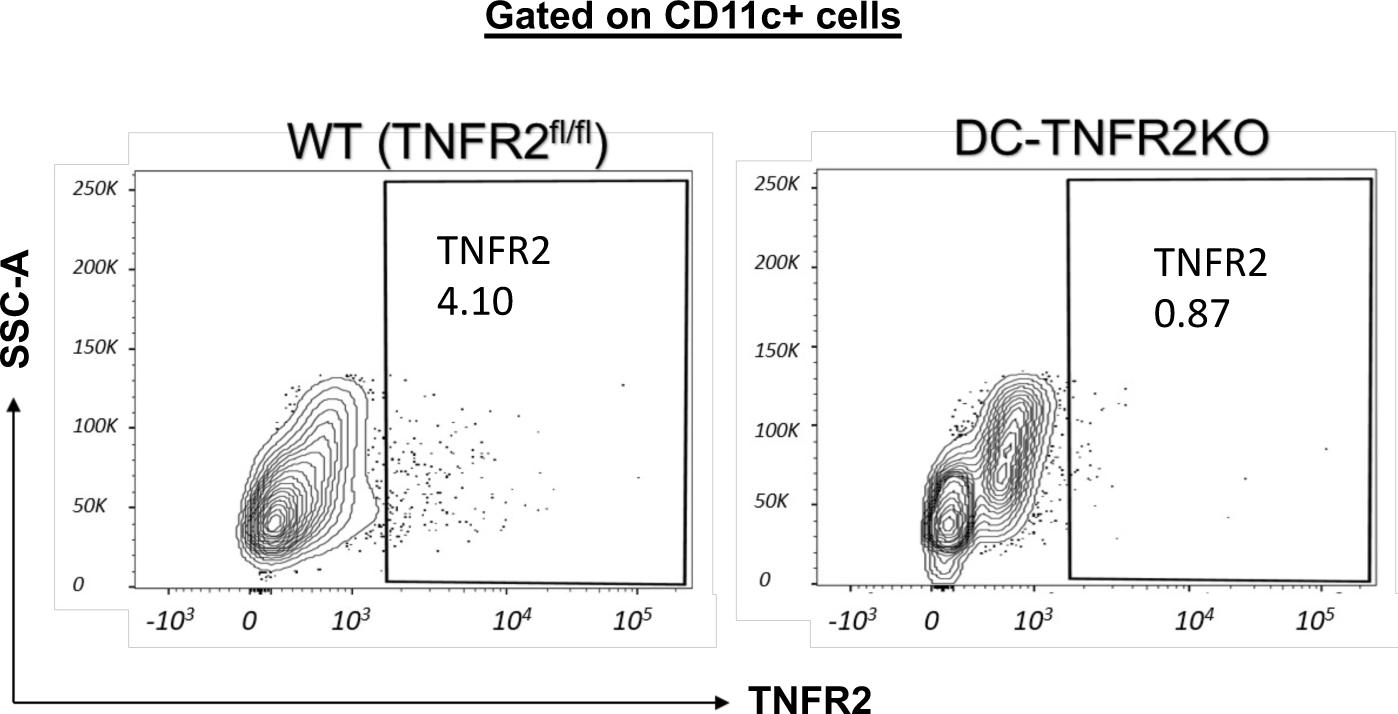
TNFR2 depletion in spleen cells isolated from DC-TNFR2KO mice. Spleen cells isolated from control (TNFR2 fl/fl) or DC-TNFR2KO mice were undergone FACS analysis and determined TNFR2 expressing CD11c+ve cells (%).

**Supplementary Figure 2.**
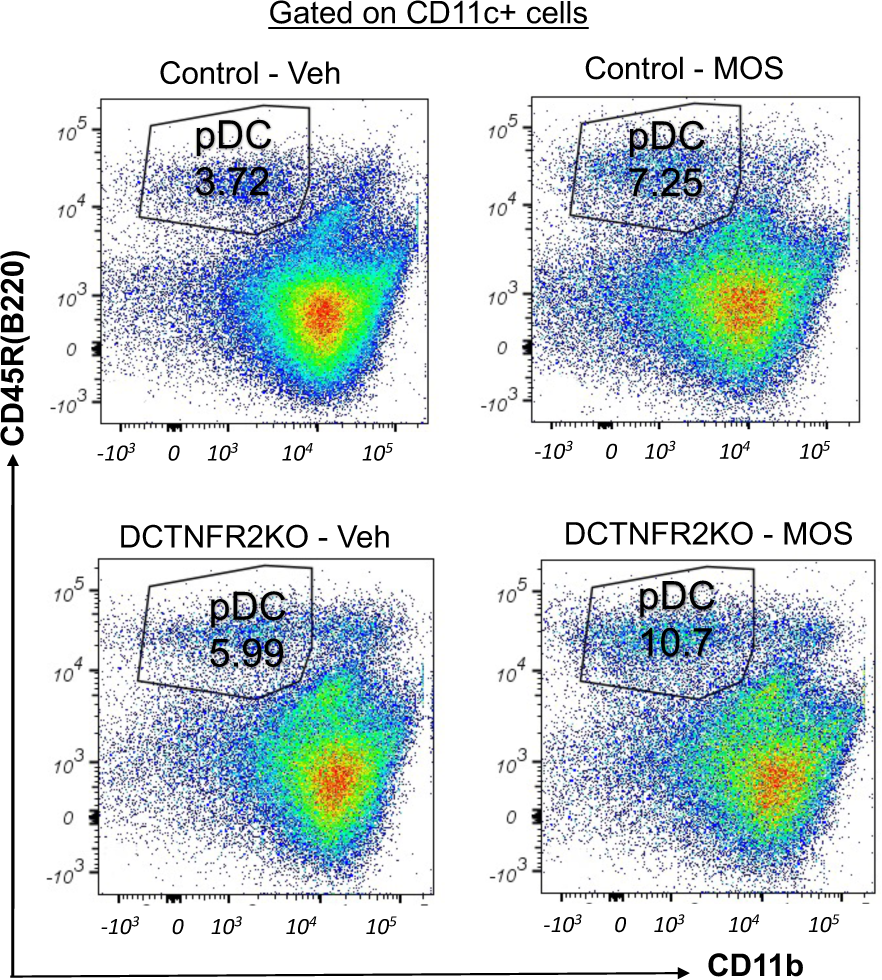
Representative FACS data showing percentage of pDC in control and DC-TNFR2KO mice ± MOS.

**Supplementary Figure 3.**
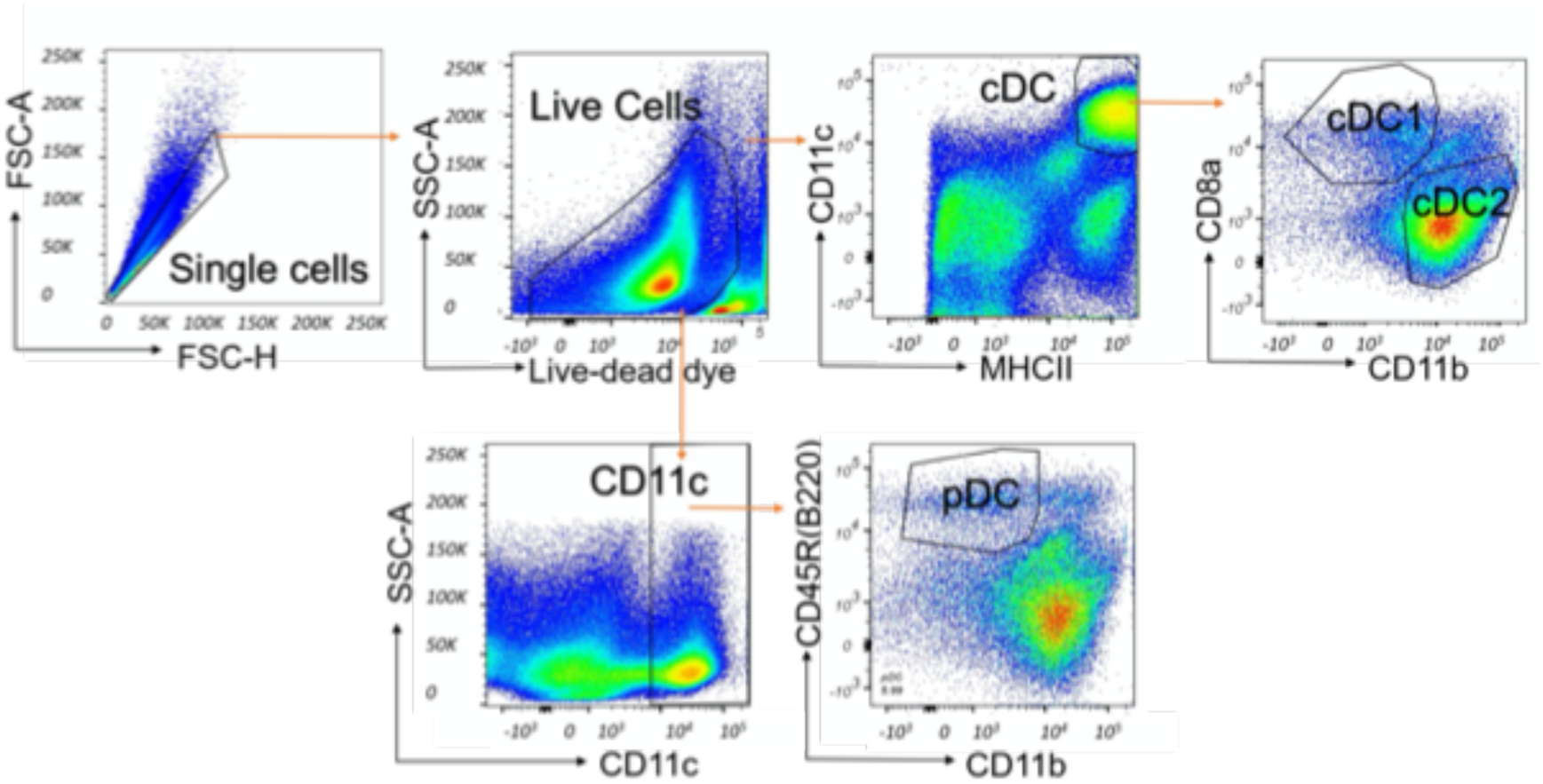
Graphical representation of the gating strategy to determine DC subsets.

